# Identification of an early-heading mutant in Indonesian native rice cultivar: ‘Gemdjah Beton’

**DOI:** 10.1101/2025.06.19.660500

**Authors:** Asanga D. Nagalla, Ryouhei Morita, Hiroyuki Ichida, Yoriko Hayashi, Yuki Shirakawa, Tadashi Sato, Yoshimichi Fukuta, Kinya Toriyama, Hiroki Saito, Yutaka Okumoto, Tomoko Abe

## Abstract

Rice is one of the world’s most widely cultivated field crops and a short-day adapted plant. It possesses a complex genetic regulatory mechanism for heading date determination. Gemdjah Beton (GB) is an Indonesian rice cultivar that takes approximately 17 weeks to flower after being transplanted into a rice field in Japan. We isolated early-heading mutant line GB-10 (flowers around two to three weeks early in the field). Under long-day (LD) conditions, GB-10 flowered approximately two weeks earlier than GB. Under short-day conditions, the heading date between GB-10 and GB plants shows no apparent difference. In agreement with the heading date, *Hd3a* and *RFT1* expressions show around a 10-time relative transcript quantity difference under LD conditions in GB-10 compared to GB. Interestingly, *Ghd7* expression was significantly reduced in GB-10, which may trigger the *Hd3a* and *RFT1* activation. A bulk segregation analysis indicated that a single nucleotide variant on chromosome 7 was genetically linked to the early heading phenotype of GB-10 lines. Taken together, we reported the successful isolation of GB-10 as an early-heading mutant.

## Introduction

Rice is one of the most widely cultivated fields of crops in the world, and almost half of the world’s population depends on it (Fukagawa and Ziska 2019). Since rice is cultivated as a field crop in wide cultivation areas, some cultivars have developed important adaptations through artificial selection to survive in particular environmental conditions. These unique phenotypic features are a product of genetic variations, and collectively, rice genetic resources play a vital role in enhancing rice yield.

Gemdjah Beton (GB) is an Indonesian rice cultivar under the ecotype Bulu. The plant has remarkable agronomic characteristics such as higher plant height, lengthier leaves, larger panicles and thick crown roots above the soil surface. This root-forming behavior, soil-surface roots (SOR), aids GB plants to adapt to anaerobic environments. The SOR in GB plants plays an important role in saline stress reduction, and the SOR is controlled by a gene known as QTL for *SOIL SURFACE ROOTING 1*(*qSOR1*) (Kitomi *et al*. 2020). In addition, GB plants can survive in phosphorous-deficient conditions due to SOR facilitating phosphorous uptake (Oo *et al*. 2021). Given these beneficial characteristics, GB is considered a reasonable breeding material for lowland rice cultivation. However, its extended vegetative period is one of the GB’s major drawbacks when selecting it as a parental line.

A complex genetic regulatory network determines rice heading date and heading date phenotype is an important adaptation to the day length variation. Rice is typically a short-day plant (Izawa *et al*. 2000), and it delays heading under Long-day (LD) conditions by promoting floral repressors. One of the key LD-dependent floral repressor is *Grain number, Plant height and heading date7* (*Ghd7*) (Xue *et al*. 2008). *Ghd7* mainly accumulates light signals from phytochromes (Osugi *et al*. 2011, Zheng *et al*. 2019, Nagalla *et al*. 2021). *Early heading date 1* (*Ehd1*) is a floral promoter in rice for both LD and short-day (SD) conditions (Doi *et al*. 2004) and *Ghd7* directly regulates it. Downstream of *Ehd1*, there are main rice florigens called *Heading date 3a* (*Hd3a*) and *Rice FT-like 1* (*RFT1*), which are the mobile flowering activators (Kojima *et al*. 2002, Komiya *et al*. 2009). In addition to this monocot-specific pathway: *Ghd7*-*Ehd1*—*Hd3a/RFT1*, rice possesses *Heading date 1* (*Hd1*), known as a homolog of Arabidopsis *CONSTANS* (*CO*) (Yano *et al*. 2000). The *CO* has been identified as a strong LD-specific floral promoter in Arabidopsis (Putterill *et al*. 1995). However, its function has diversified in rice since *Hd1* advances flowering under SD conditions by promoting floral induces while it represses the flowering under LD conditions (Yano *et al*. 2000). Under LD’s daytime conditions, a post-translational interaction between *Ghd7* and *Hd1* has been reported. The Ghd7-Hd1 complex binds to the *Ehd1* promoter and represses its expression. Under SD conditions, *Hd1* alone promotes *Ehd1* expression, thus inducing flowering (Nemoto *et al*. 2016). The activities of circadian and flowering-time regulatory genes are key in determining rice plants’ vegetative to reproductive phase transition.

In the present study, we isolated an early-heading mutant line, GB-10, in the GB background. The early heading phenotype was consistent in controlled LD and natural-day length field (ND) conditions. Since the GB-10 induces heading under LD conditions, we investigated the expression of the key flowering-time regulatory gene. A population analysis shows that the mutation in the unknown gene possesses an incomplete dominance. A resequencing-based bulk segregation analysis shows that a single nucleotide variant (SNV) on an intergenic region of chromosome (Chr.) 7 is genetically linked to the early heading phenotype.

## Material and Methods

### Plant material and growth conditions

Dry rice seeds of Gemdjah Beton (GB) (*Oryza sativa* L. ecotype Bulu) were irradiated by C-ion beams (12C^6+^ ions, 135 MeV nucleon^−1^) at the dose of 125 Gy in the RI-beam factory (RIKEN, Saitama, Japan). Isolation of the early heading mutant was done as described by Hanzawa *et al*. (2014). Plants were grown in paddy fields at Experimental Farm Station, Tohoku University, Kashimadai, Osaki, Miyagi, Japan.

### Measuring days to heading

Heading date observation was conducted using GB-10 and GB plants under controlled SD, LD and ND conditions. Rice plants were grown under ND conditions in paddy fields at Experimental Farm Station Graduate School of Life Science Tohoku University, in Kashimadai (Miyagi, Japan). The SD condition was 10 hr light (28 ℃)/14 hr dark (24 ℃), and the LD condition was 14.5 hr light (28 ℃) /and 9.5 hr dark (24 ℃). Germinated seedlings were grown under SD or LD conditions and transplanted to pots after three weeks under the same conditions. The date when the first panicle emerged was recorded as the heading date.

### Gene expression analysis

The fully emerged uppermost leaves of plants 21, 49, and 90 days after sowing (DAS) were harvested 2 hr after light. Leaves from three plants were mixed as a sample. As biological replicates, three or four samples were used for Real-Time Quantitative Reverse Transcription PCR (qRT-PCR). Total RNA was extracted from rice leaves using TRIZOL reagent (Invitrogen, ThermoFisher Scientific, Waltham, MA, USA) according to the manufacturer’s instructions and treated with DNase I: TURBO DNA-free™ Kit (Invitrogen). The cDNA was synthesized using 4 μg of total RNA using ReverTra Ace qPCR RT master mix (TOYOBO, Tokyo, Japan). Real-time quantitative RT-PCR was performed with the TaqMan fast universal PCR master mix (Applied Biosystems) or THUNDERBIRDTM SYBR qPCR Mix (only the data in Fig. 5 was generated by SYBR) (TOYOBO, Tokyo, Japan) on a LightCycler 480 II (Roche, Basel, Switzerland) according to the manufacturer’s instructions. Gene-specific primers and TaqMan probe sequences are listed in Supplemental Table 1. A rice ubiquitin gene (*Os02g0161900*) was used for normalization. Normalized data were logarithmically transformed (log10) in figures considering expression dynamics and data fluctuations of related genes.

### A resequencing-based bulk segregation analysis

The F_3_ population was grown in paddy fields in Kashimadai expecting the segregation of the heading-date phenotype. Early-heading type and normal-heading type F_3_ plants were selected, and seeds were collected from each plant. Six lines of the early-heading and six of the normal-heading F_4_ plants were used for whole genome sequencing. Subsequently, equal amounts of leaf tissue were collected from five F_4_ plants of one normal-heading line and seven plants, each of the other lines. Genomic DNA was extracted from the leaf tissue using a DNeasy plant mini kit (QIAGEN, Venlo, Netherlands), with IDTE 1×TE solution pH 8.0 (Integrated DNA Technologies, Iowa, USA) instead of the AE buffer provided in the kit for the DNA elution step. Equal amounts of extracted DNA were mixed to create “bulked DNA of normal-heading type” and “bulked DNA of early-heading type”. Each bulked DNA library was prepared using an MGIEasy PCR-Free DNA library prep kit (MGI Tech, Shenzhen, China). Whole genome sequencing was performed using the MGI DNBSEQ-G400 instrument (MGI Tech) in paired-end, 2 × 150-bp mode. Bioinformatics analysis used the mutation analysis pipeline (Ichida *et al*. 2019). In the mutation analysis pipeline, mutations were detected using GATK v3 (Mckenna *et al*. 2010), BcfTools (Li, 2011), Pindel (Ye *et al*. 2009), Delly (Rausch *et al*. 2012), and Manta (Chen *et al*. 2016). The genome sequence of the japonica cultivar Nipponbare (Os-Nipponbare-Reference-IRGSP-1.0, Kawahara *et al*. 2013) was used as a reference genome sequence. The resulting candidate mutations were visually confirmed using the Integrative Genomics Viewer software (IGV; Robinson *et al*. 2011). Nucleotide sequence data files are available in the NCBI Sequenced Read Archive under the accession number PRJNA1225329 (https://www.ncbi.nlm.nih.gov/sra/PRJNA1225329).

### PCR and Sanger-sequence for determining the genotype of the SNV on chromosome 7

We grew 50 F_3_ plants as a segregation population. Genomic DNA was extracted from each F_3_ plant using QuickExtract^TM^ Plant DNA extraction solution (LGC, London, UK). PCR was performed in a total volume of 20 µl, containing 1 µl of genomic DNA, 10 µl of 2 × Gflex PCR buffer (Takara Bio, Shiga, Japan), 4 pmol of each primer, and 0.5 Unit of Tks Gflex DNA polymerase (Takara Bio). The PCR was carried out with an initial denaturation step at 94℃ for 1 min, then 35 cycles of 98℃ for 10 s, 60℃ for 15 s, and 68℃ for 45 s. Sequences of each primer were as follows: F: 5’-AGACGGCCAACAGAGGGAATG-3’. R: 5’-CCGATTACTCCTGCCGATCT-3’. Sanger sequencing was performed to determine the genotype of the SNV using BigDye^TM^ Terminator v 3.1 Cycle sequencing kit (Thermo Fisher Scientific, Massachusetts, USA) and ABI 3730xl Genetic Analyzer (Thermo Fisher Scientific).

## Results

### A Gemdjah Beton mutant, GB-10, shows an early heading date under Long-day conditions

We observed early heading phenotype in GB-10 in ND conditions for two years (Table 1). The heading date of GB-10 was about 15-21 days earlier than that of GB. For characterization of early heading GB-10, we grew GB and GB-10 plants under SD and LD conditions (Fig. 1). The GB showed a clear photoperiod response, flowering 38 days later than SD under LD conditions. GB-10 flowered around 16 days earlier than that of GB under LD conditions. However, under SD conditions, GB-10’s flowering tended to be delayed, but there was no difference in flowering time between GB and GB-10 (Table 1).

**Fig. 1.**
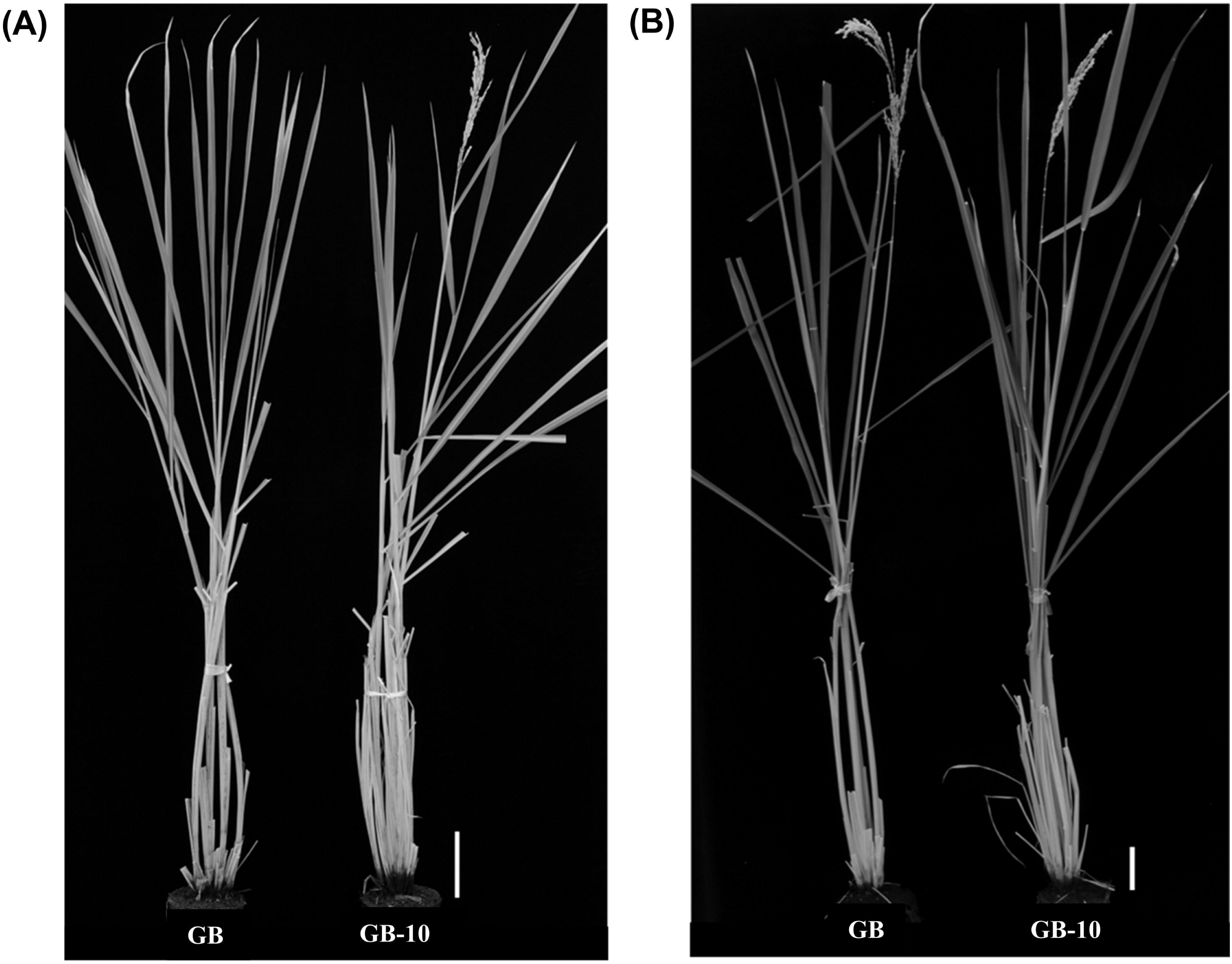
The heading dates of the GB and GB-10 mutant under LD and SD conditions. **A** GB and GB-10 plants grown under LD conditions, photos were taken when the GB-10 plants flowered around 150 DAS. **B** GB and GB-10 plants grow under SD conditions, photos were taken when the GB-10 plants flowered around 128 DAS. The experiment was performed two times and observed the same results. Bar size = 10cm. Figure Size: 12.33 cm (H) × 15.59 cm (W)

**Table 1.**
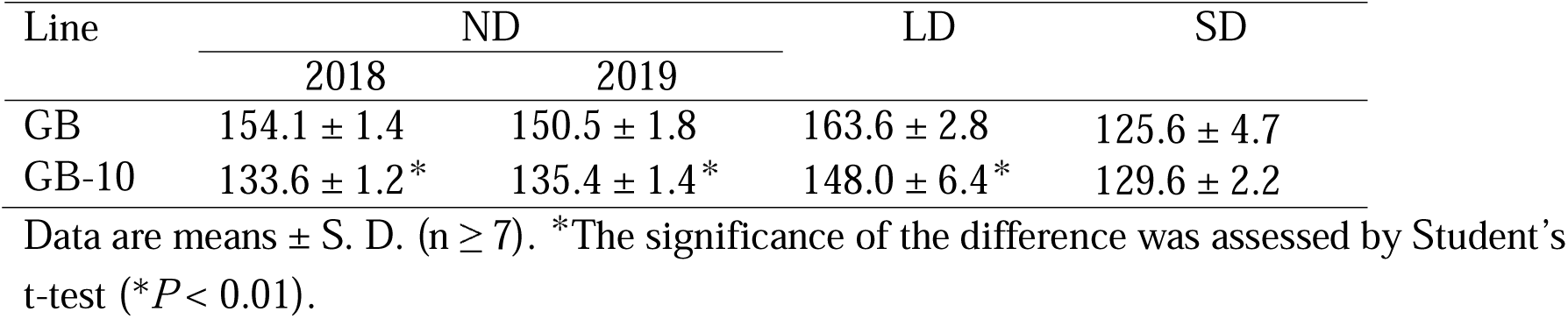
The heading dates of the GB and GB-10 under various daylength conditions.

### Florigen expression is consistent with early heading date phenotype in GB-10

Since the GB-10 shows a clear induction of heading under LD conditions, we investigated the expression of the key circadian clock and flowering time gene expression. Based on the days to heading data, we observed that GB plants required more than 120 days for heading in both LD and SD conditions (Table 1). Therefore, gene expression was analyzed in the 21 DAS seedlings and 49 DAS plant stages (Fig. 2). *Ghd7* is a key repressor under LD conditions (Xue *et al*. 2008; Itoh *et al*. 2010). Its expression has reduced in the GB-10 compared to GB in both 21 and 49 DAS plants. This de-repressive effect promoted rice florigen coding genes *Hd3a* and *RFT1* (Kojima *et al*. 2002) (Fig. 2). The consistent florigen expression promotion from 21 DAS to 49 DAS in GB-10 may result in an early heading compared to GB under LD conditions. Under SD conditions, GB and GB-10 plants did not show a significant difference in heading date (Table 1). The *Hd1* is an upstream photoperiodic flowering regulator under SD conditions (Yano *et al*. 2000). Its expression showed no significant difference between GB and GB-10 in 21 and 49 DAS plants. These *Hd1* transcript levels have affected downstream genes, and it might lead to the comparable heading date between GB and GB-10. The *Ehd1*, a floral promoter under both SD and LD conditions (Doi *et al*. 2004). Consistently with comparable heading dates in SD conditions, *Hd3a* and *RFT1* quantity differences were low in SD conditions compared to LD conditions in GB and GB-10 lines (Fig. 2). Specifically, GB and GB-10 in 49 DAS, *RFT1* expression has resulted in equal quantities providing evidence to result comparable heading dates. Meanwhile, the florigen expression in GB and GB-10 under LD conditions showed a clear difference (Fig. 2), indicating the genetic effect in regulating photoperiodic flowering induced by the mutations that occurred in GB-10.

**Fig. 2.**
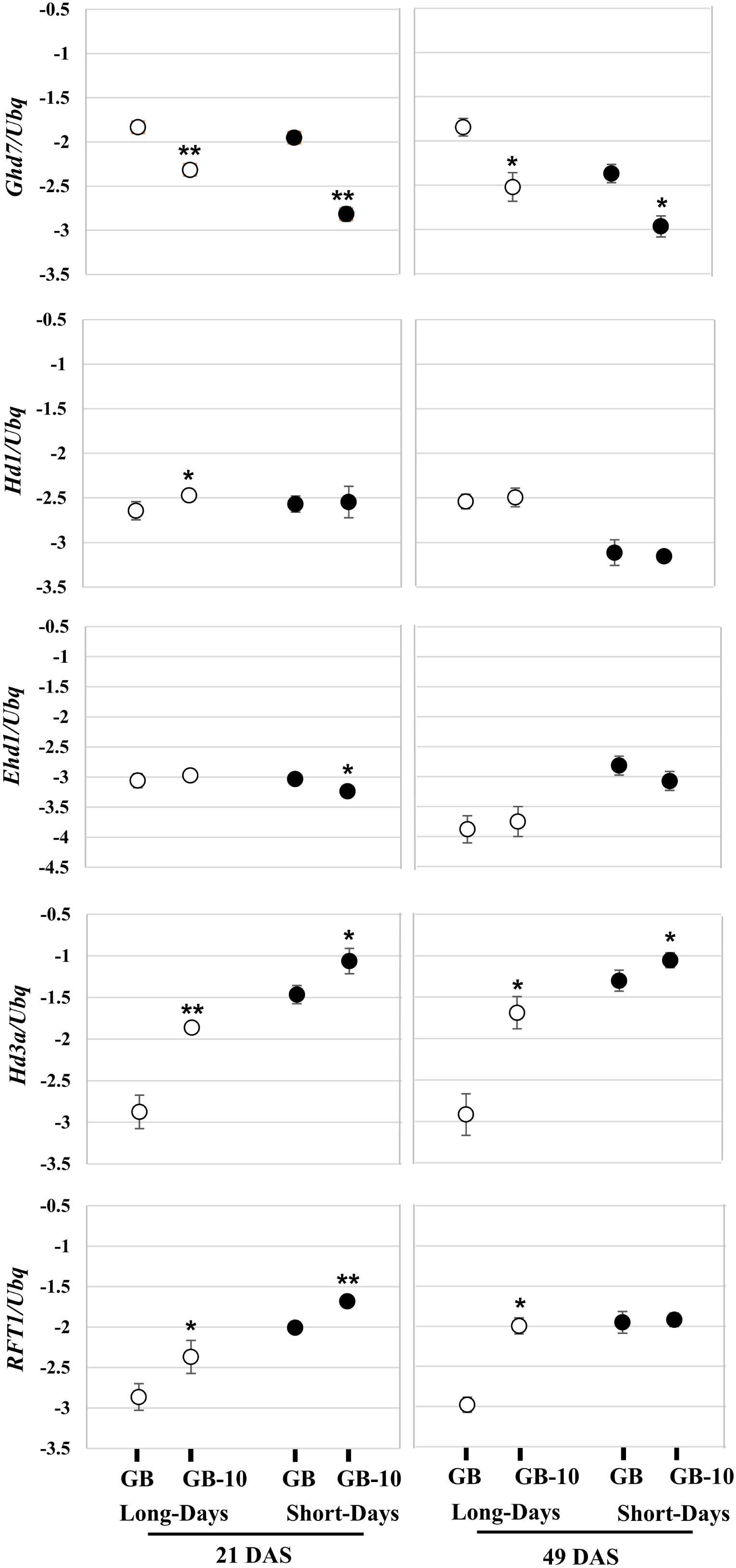
Relative gene expression of *Ghd7*, *Hd1*, *Ehd1*, *Hd3a* and *RFT1*. 21days old (left) and 49 days old (right) leaf blade samples were harvested 2hr after dawn. Data are means ± S. D. (n = 3 or 4 biological replicates); and the significance of the difference was assessed by Student’s t-test (\*\**P* < 0.01; \**P* < 0.05). Relative gene expression was shown in the logarithmic Y-axis. Rice ubiquitin gene (*Os02g0161900*) was used for normalization. Figure Size: 26.2 cm (H) × 12.4 cm (W)

### Inheritance pattern of the early-heading phenotype of GB-10

We generated an F_2_ population derived from the cross between GB-10 and GB. We grew it in a paddy field to investigate how the early-heading phenotype in GB-10 was inherited. The distribution of heading dates in the F_2_ individuals was continuous (Fig. 3). When we classified the F_2_ individuals into early-heading type, normal type, and intermediate type based on the heading dates of the parents, the ratio of early-heading, intermediate, and normal types fitted a 1:2:1 ratio (25:53:29, chi-square score = 0.31, *p* > 0.05). These results indicate that a single locus confers the early-heading phenotype of GB-10 and exhibits incomplete dominance.

**Fig. 3.**
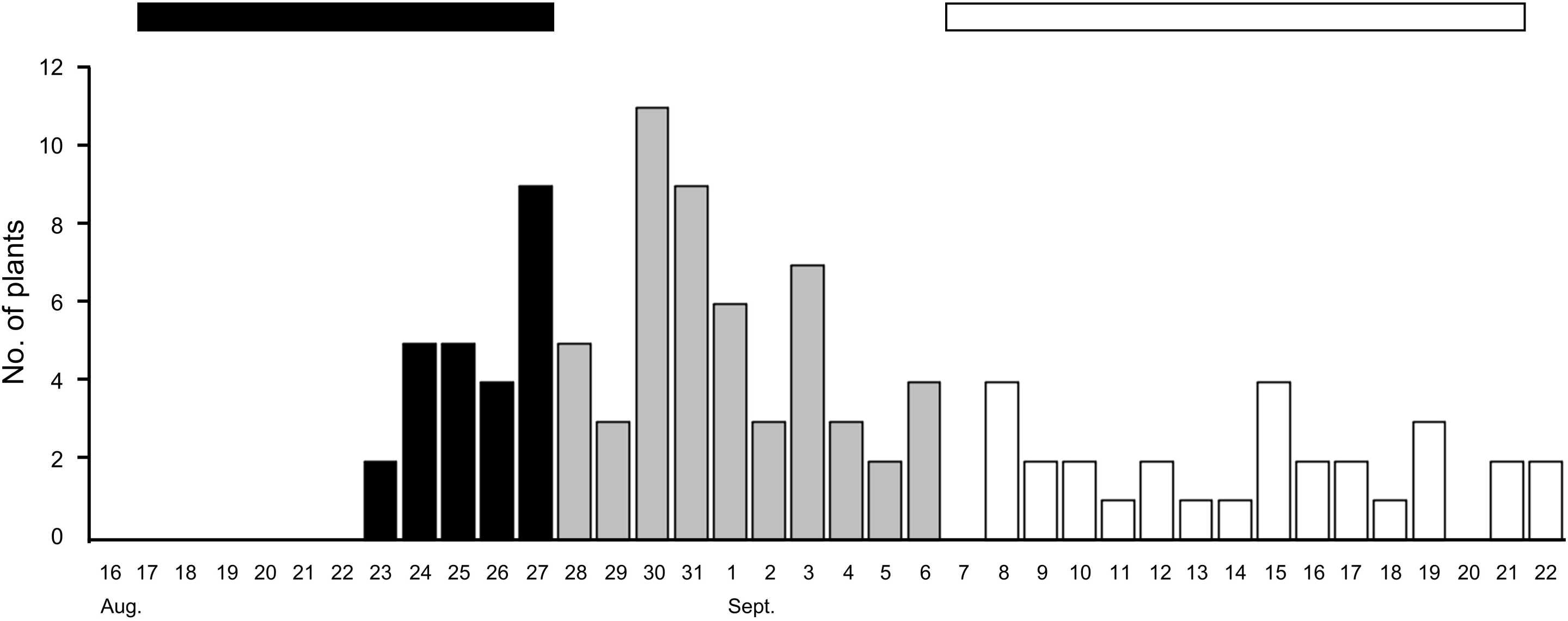
Distribution of heading days in the F_2_ population derived from the cross between GB and GB-10. Black and white bars indicate the ranges of heading days of GB and GB-10, respectively. Figure Size: 10.8 cm (H) × 24.6 cm (W)

*An SNV on* chr. *7 is genetically linked with early heading in GB-10*

We performed a resequencing-based bulk segregation analysis (Li and Xu, 2022) to identify the gene or region responsible for the early-heading phenotype using the F_4_ plants. We analyzed both “bulked DNA of early-heading type” and “bulked DNA of normal-heading type”. We obtained the sequence data for a total of 120.0 and 106.6 million reads, with 34.7 and 30.9 billion bases aligned for “bulked DNA of normal-heading type” and “bulked DNA of early-heading type”, respectively (Supplemental Table 2). The average coverages were 93.0× and 82.7×, respectively. Sixty-three mutations were detected in the “bulked DNA of early-heading type” sample (Supplemental Table 3). We checked the mutation output from the mutation analysis pipeline using IGV and confirmed that 50% of the detected mutations were positive (Supplemental Table 4). We detected 63 SNVs in total. In bulk segregation analysis, only mutations occurring within the responsible gene or its vicinity exhibit a higher SNP index (with the maximum SNP index being 1.0, Abe *et al*. 2012). Of the 63 detected SNVs, one SNV (G to A) at position 10559536 on chr. 7 exhibited the highest SNP index of 0.9, while the SNP index of the remaining 62 SNVs was 0.6 or lower. These findings indicated that the SNV at chr.7: 10559536 was genetically linked to the early-heading phenotype of GB-10. In addition, there were no mutations in the *qSOR1* gene region (Locus ID: *Os07g0614400*), indicating that GB-10 is useful as a parental line for the development of a new variety with a soil-surface rooting phenotype (Data not shown).

The position where the SNV occurred (chr.7: 10559536) had been identified as an intergenic region by SnpEff (Cingolani *et al*. 2012) software (Fig. 4). The distance from the SNV to known flowering time regulatory or circadian clock regulatory genes on the chr. 7 was quite far. Next, we investigated whether this SNV affects the expression levels of the surrounding four genes (*Os07g0278400*, *Os07g0278866*, *Os07g0280200*, and *Os07g0280600*) identified based on IRGSP-1.0 in the RAP-DB database (Sakai *et al*. 2013). We performed qRT-PCR using the cDNA produced from total RNA from leaf blade samples (Supplemental Table 1). However, there was no significant difference among the transcript levels of the target four genes among GB and GB-10 (Fig. 5). This result indicates that SNV on chr.7 had no regulatory effects on any of the four surrounding genes, suggesting that these genes were not involved in the early-heading phenotype of GB-10.

**Fig. 4.**
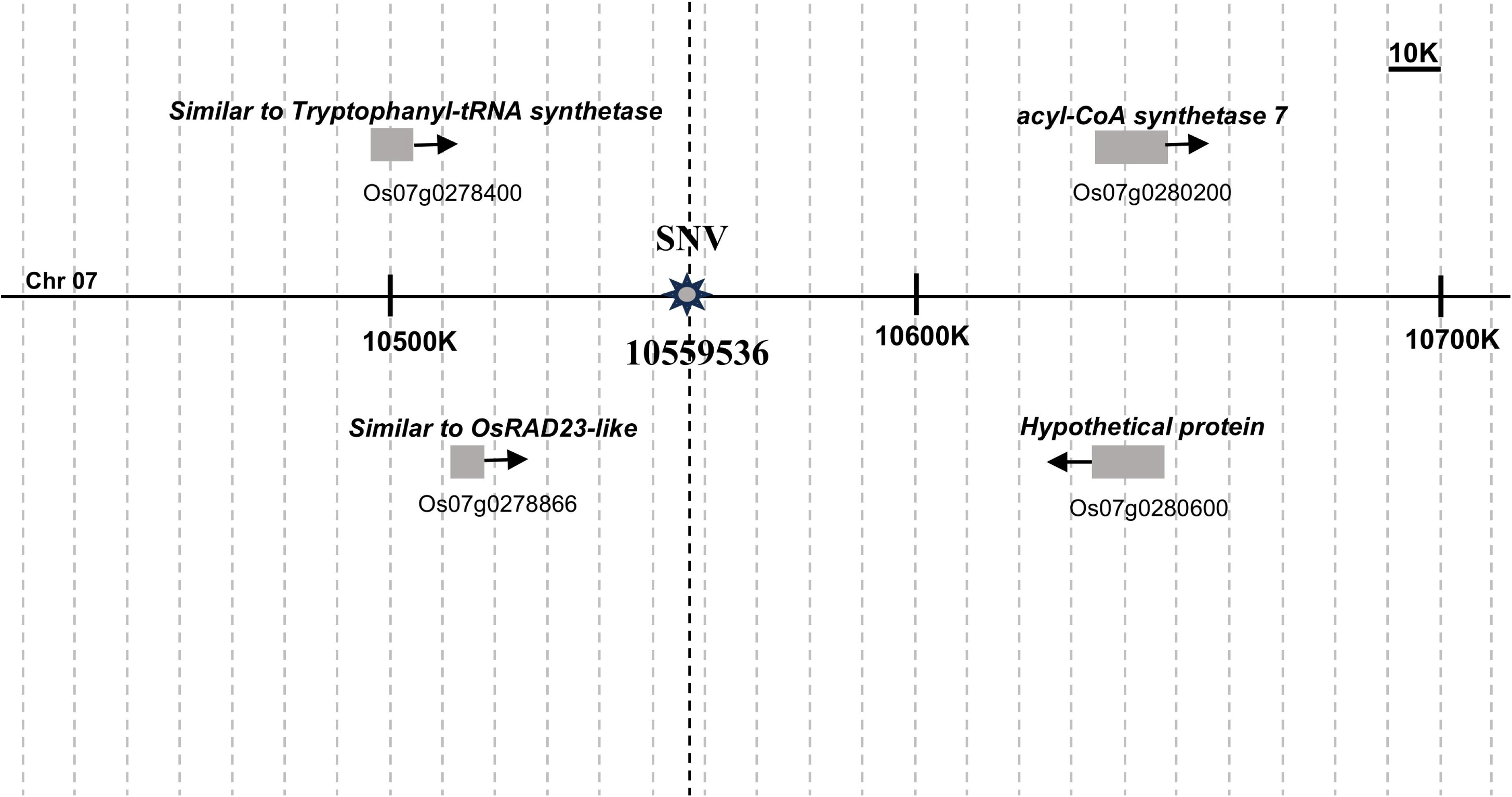
Representation of a fraction of the physical map of Chromosome 7 showing the position of the candidate SNV and four adjacent genes. Gene annotations and relative positions were based on the RAP-DB. Figure Size: 8.1 cm (H) × 15.4 cm (W)

**Fig. 5.**
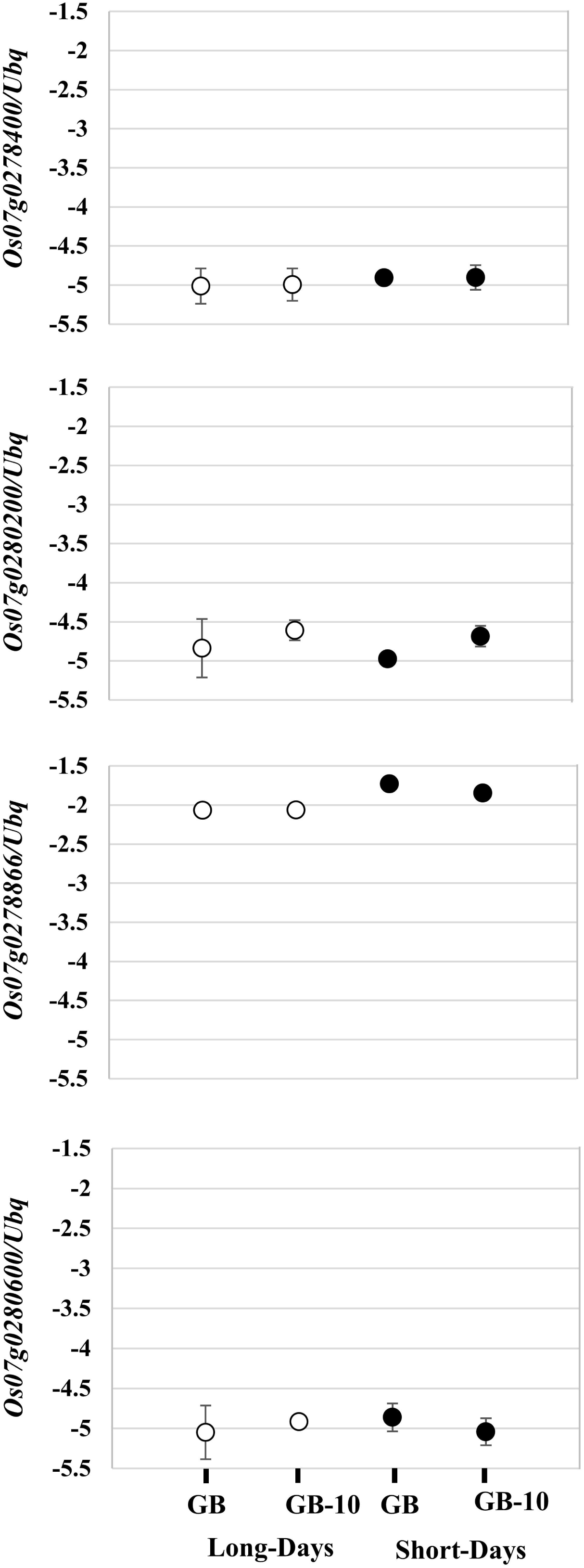
Relative gene expression of four adjacent genes *Os07g0278400, Os07g0278866, Os07g0280200*, and *Os07g0280600*. Leaf blades of seven-week-old plants of GB and GB-10 were harvested 2hr after dawn. Relative gene expression was shown in the logarithmic Y-axis and rice ubiquitin gene (*Os02g0161900*) was used for normalization. Data are means ± S. D. (n = 3 or 4 biological replicates). Figure Size: 22.7 cm (H) × 8.8 cm (W)

### The inheritance pattern of the early-heading phenotype of GB-10

To confirm the inheritance pattern of the early-heading phenotype of GB-10, an F_3_ segregation population derived from a single F_2_ plant that retained the heterozygous SNV on chr. 7 was grown under LD conditions. Genotyping analysis of the SNV of each F_3_ plant revealed that 12 plants had homozygous SNV, 24 had heterozygous SNV, and 14 did not possess SNV. The heading dates (DTH) for the F_3_ plants with homozygous SNV were 136.7 ± 3.2 (means ± S.D.). In contrast, the DTH of plants without the SNV was 151.4 ± 7.4, further confirming that the SNV was genetically linked to the early-heading phenotype. The DTH for F_3_ plants with the heterozygous SNV was 143.9 ± 5.1, showing an intermediate phenotype between the plants having the homozygous SNV and those without the SNV. These results demonstrate that the early-heading phenotype of GB-10 exhibits incomplete dominance, as in the F_2_ population.

## Discussion

Gemdjah Beton is a native Indonesian rice cultivar. It possesses several agronomically important phenotypic features such as higher plant height, column length, lengthier leaf blades (Fig. 1), larger panicle and strong surface root system. Based on these characteristics, GB plants have become a strong landrace. Therefore, breeders use GB plants as parent materials in their genetic improvement procedures (Kitomi *et al*. 2020, Oo *et al*. 2021). However, the major constraint to adapting GB as a breeding material is its lengthier vegetative phase, which spans over 150 days in ND conditions in the northern part of Japan (Table 1). Due to this extended vegetative phase, GB plants cannot produce filled grains in field conditions, as the ambient temperature drops within the grain-filling stage around late October. GB-10 plants flower around 15-21 days earlier than GB under ND conditions, and it is able to obtain the filled gains.

The main determinants for rice flowering time or heading date are key environmental stimuli such as day length conditions, circadian clock and ambient temperature. They are recognized by a set of genes, and those genes are responsible for phenotypic variations in flowering time. The flowering time and circadian clock gene expression in GB and GB-10 correspond to the early heading date of GB-10. However, these observations were made with Nipponbare plants, where flowering time is usually around 100 DAS under LD and 60 DAS under SD conditions. GB plants show prolonged vegetative periods (Table 1 and Fig. 1), and 21 DAS seedling’s florigen expression results may be insufficient as representative data. Therefore, we checked another data point at the later vegetative stage to confirm florigen expression levels. Both 21 and 49 DAS plants resulted in almost consistent florigen gene expression results. Because the difference in the *Hd3a* and *RFT1* expression is greater in the 49 DAS plants, 49 DAS plants were suitable for explaining the flowering time gene expression and heading date phenotypes in GB and GB-10 plants.

The rice flowering time regulatory pathways have been illustrated based on the evidence using *O. sativa* L. ‘Nipponbare’ plants. Therefore, we measured the transcript levels of the same key flowering time genes in Nipponbare leaves and compared them with GB plants (Supplemental Fig. 1). Previous reports indicate that early flowering plants (mutants or NILs) consistently show higher *Hd3a* and *RFT1* expression levels throughout their vegetative periods compared to their wild-type plants (Kojima *et al*. 2002, Nemoto *et al*. 2016). It was interesting to observe that Nipponbare resulted in early heading compared to GB under LD conditions. However, its *Hd3a* and *RFT1* expression in the 21 DAS seedlings were significantly lower than that of GB. This data represents the genetic and allelic distance of heading date and circadian-clock regulatory genes between Nipponbare and GB.

Under LD conditions, the functional allele of *Ghd7* in Nippponbare represses the *Ehd1* and subsequently represses the *Hd3a* and *RFT1* (Xue *et al*. 2008, Itoh and Izawa 2013, Nemoto *et al*. 2016, Nagalla *et al*. 2021). However, this conventional *Ehd1* repression was not observed in GB plants in 21 and 49 DAS plants (Fig. 2). The reduced amount of *Ghd7* transcripts in GB-10 does not result in a clearly elevated *Ehd1* transcript. However, a significant *Hd3a* and *RFT1* expression was observed. Interestingly, we observed a significant *Ehd1* induction in GB-10 plants in 90 DAS (Supplemental Fig. 2), indicating that *Ehd1* induction occurred around the latter vegetative periods in the GB variety. Notably, 90 DAS GB-10 plants do not show a clear *RFT1* transcript difference between GB, but these plants show an early-heading phenotype (Supplemental Fig. 2). Therefore, under LD conditions, 90 DAS plants’ *Ghd7* reduction may directly affect the *Ehd1* de-repression, which results in *Hd3a* induction. The *RFT1* induction was not observed, probably due to the developmental stage’s effect on the *RFT1* expression. Altogether, *Ghd7*, *Hd1*, *Hd3a*, and *RFT1* expressions are generally consistent with the heading date phenotype of GB plants and their role in conventional heading date regulatory pathway is consistent with Nipponbare as we did not find any allelic differences or polymorphisms in GB compared to Nipponbare.

The SNV was positioned on the intergenic region based on the short-read sequences mapped to the Nipponbare reference genome (Fig. 4). None of the surrounding genes were related to flowering time control or circadian clock regulation. A comparable gene expression was observed with GB and GB-10 plants for four surrounding genes tested, indicating the SNV did not affect their expression (Fig. 5). Therefore, those genes should not be responsible for the early flowering phenotype of GB-10. However, the linkage analysis results showed that SNV and the early flowering phenotype are tightly linked, and the position of the responsible gene should be close to the SNV. Therefore, the SNV on the chr. 07: 10559536 can be used as a DNA marker to identify the responsible gene for early flowering phenotype. The SNV position remains intergenic based on the IRGSP-1.0 to date (Assessed 2024/11/04). We did the short-read sequencing and mapped it into the reference genome IRGSP-1.0. This approach may hide some important genes that occurred exclusively in the Gemdjah Beton genome. Creating a Gemdjah Beton reference genome can be suggested as a future perspective that would be beneficial in identifying the unknown responsible gene. Notably, the responsible gene proved to have an effect upstream of *Ghd7* and possesses an important potential on the rice flowering time regulatory pathway.

## Supporting information

Supplemental Table 1, and will be used for the link to the file on the preprint site

## Author contributions

Conceptualized by T.A. and T.A, R.M. and A.D.N designed the experiments. T.A., Y.H., Y.S., T.S., Y.F., K.T., H.S., Y.O., R.M. and A.D.N performed experiments and data analysis. H.I. performed NGS analysis and mutation calling pipeline. A.D.N, R.M. and T.A. wrote the manuscript. T.A. did project administration and funding acquisition. All authors have read and agreed to the published version of the manuscript.

## Acknowledgments

We thank the RIKEN Nishina Center and the Center for Nuclear Study, the University of Tokyo for operating RIBF for performing the ion-beam irradiations. We are grateful for the technical help the Support Unit for Bio-Material Analysis, RIKEN CBS Research Resources Division provides regarding Sanger sequencing. The bioinformatics analysis was performed using the HOKUSAI-BigWaterfall supercomputing system (RIKEN, Saitama, Japan) under project numbers Q22208 and Q23443. This project was supported by the Cross-ministerial Strategic Innovation Promotion Program (SIP) “Technologies for creating next-generation agriculture, forestry and fisheries” (funding agency: Bio-oriented Technology Research Advancement Institution, NARO).

