## Supplemental Table 1, and will be used for the link to the file on the preprint site for "Identification of an early-heading mutant in Indonesian native rice cultivar: ‘Gemdjah Beton’"

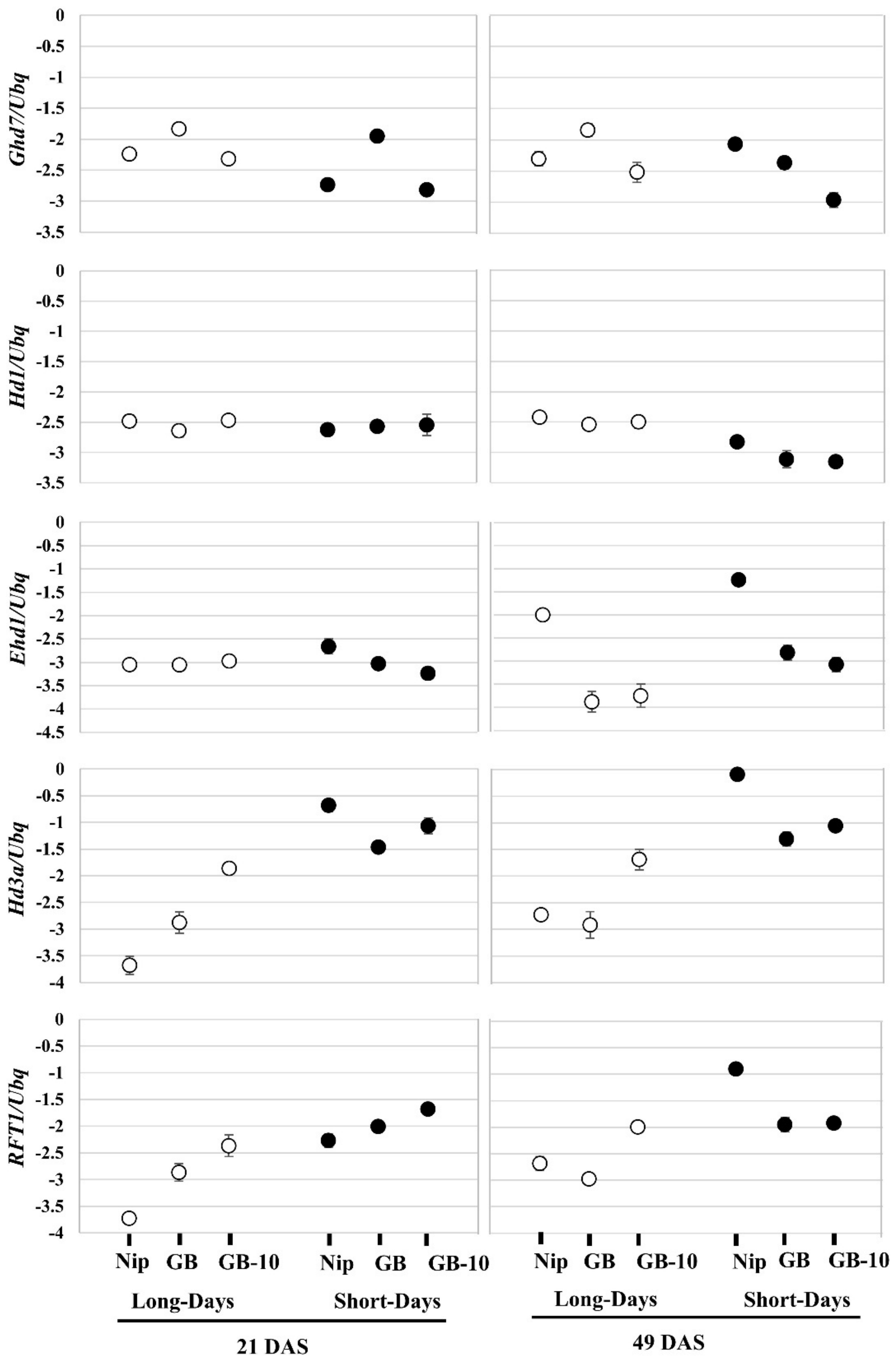


**Supplemental Fig. 1.**

**Relative gene expression of *Ghd7*, *Hd1*, *Ehd1*, *Hd3a* and *RFT1*. of the cv. Nipponbare, GB and GB-10 mutant under LD and SD conditions**. 21days old (left) and 49 days old (right) leaf blade samples were harvested 2hr after dawn. The significance of the difference was assessed by Student's t-test (***P* < 0.01). The relative gene expression was shown in the logarithmic Y-axis.

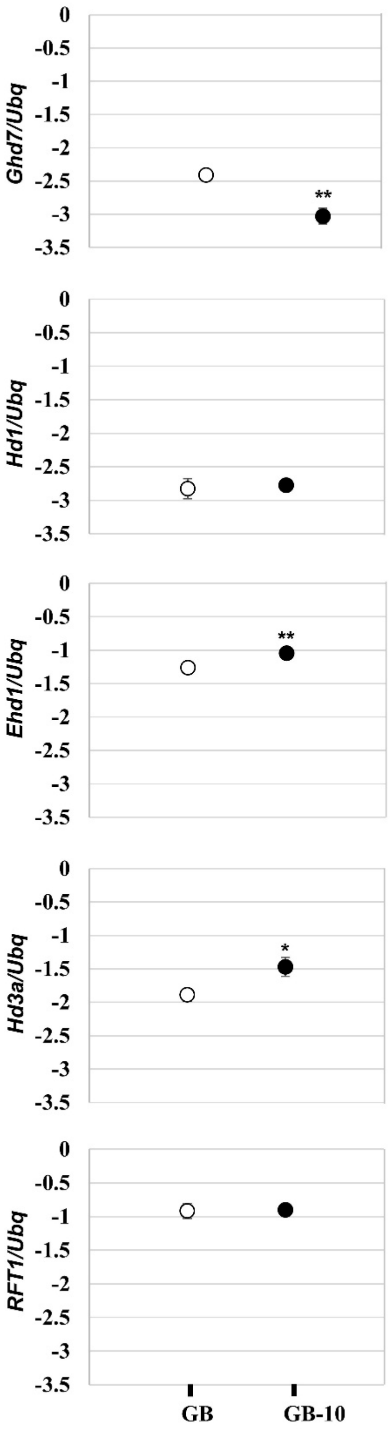


**Supplemental Fig. 2.**

**Relative gene expression of *Ghd7*, *Hd1*, *Ehd1*, *Hd3a* and *RFT1*. of the GB and GB-10 mutant under LD conditions.** 90 DAS plant leaf blade samples were harvested 2hr after dawn. Data are means ± S. D. (n = 3 or 4 biological replicates). Relative gene expression was shown in the logarithmic Y-axis.

**Supplemental Table 1.** Primers and probe sequences used in this study

| **Primer** | **Primer sequence （5'-3'）** | **Taqman Probe** | **Probe sequence** |
|---|---|---|---|
| *Ubq*_RT_fw | GAGCCTCTGTTCGTCAAGTA | *Ubq*_Probe | TTGTGGTGCTGATGTCTACTTGTGTC |
| *Ubq*_RT_Rw | ACTCGATGGTCCATTAAACC | | |
| *Ghd7*_RT_fw | GTACGCGTCCAGAAAAGCT | *Ghd7*_Probe | TGCCGAGATGAGGCCCCGA |
| *Ghd7*_RT_Rw | TTGGCGAAGCGACCTCTC | | |
| *Hd1*_RT_fw | AGCAGCATAGTGGTTATGGAGTTG | *Hd1*_Probe | ACACAGATTCCATCAGCAACAGCATATCTT |
| *Hd1*_RT_Rw | CACCGTGCTGTCTGGTACTATAC | | |
| *Ehd1*_RT_fw | GAGGATCGAAGAGCTGAGCA | *Ehd1*_Probe | CATTTGGCAGCACATATTCCGAAAGCA |
| *Ehd1*_RT_Rw | AGGATGACCGGGTTTTTCGA | | |
| *Hd3a*_RT_fw | TCTACTTCAACTGCCAGCGC | *Hd3a*_Probe | TCCCGATCGATCTGCTGCATGC |
| *Hd3a*_RT_Rw | TTCAATTGTCTGAACCTGCAATGT | | |
| *RFT1*_RT_fw | CAGAACTTCAGCACCAGGAACTT | *RFT1*_probe | AGCTCTACAACCTCGGCTCGCCG |
| *RFT1*_RT_Rw | TCGCGCTGGCAGTTGA | | |
| *Os07g0278400*_RT_fw | GAAAAGTTTATGTGGAAGAACTTGACTATT | | |
| *Os07g0278400*_RT_Rw | TTTTTCAATGTCAAATCCACAGGTGATAA | | |
| *Os07g0278866*_RT_fw | CTAGATTTGGCTGTTAACTGTGTTGTAAAT | | |
| *Os07g0278866*_RT_Rw | TTTATCATACAATCTCAGATTCAAACCAACA | | |
| *Os07g0280200*_RT_fw | CAGGTGGCATATCAAGGATGATCAG | | |
| *Os07g0280200*_RT_Rw | AACAATCTTTACCTCGACATTCTCAG | | |
| *Os07g0280600*_RT_fw | ACGAACAAGAACCCCATGGAG | | |
| *Os07g0280600*_RT_Rw | GGAGCTTCATCGCCATGGTG | | |

**Supplemental Table 2.** Summary of WGS in $F_4$ plants.

| Sample name | Normal- heading bulk | Early-heading bulk |
|---|---|---|
| Total reads | 120037225 | 106614296 |
| Total reads aligned | 118198119 | 105101608 |
| Reads aligned (%) | 98.5 | 98.6 |
| Total bases aligned | 34719858820 | 30871332241 |
| Average coverage | 93.0 | 82.7 |

**Supplemental Table 3.** List of SNVs in the GB-10 mutant.

| Mutation ID | Chr. | Position | Ref | Alt | Type of mutation | Effect | SNP index |
|---|---|---|---|---|---|---|---|
| 1 | 1 | 7903845 | C | G | SNV | missense_variant | 0.4 |
| 2 | 1 | 16087524 | A | G | SNV | missense_variant | 0.5 |
| 3 | 1 | 18909974 | C | G | SNV | intergenic_region | 0.3 |
| 4 | 1 | 19810007 | G | A | SNV | upstream_gene_variant | 0.3 |
| 5 | 1 | 22732057 | C | G | SNV | synonymous_variant | 0.3 |
| 6 | 1 | 40712713 | C | G | SNV | missense_variant | 0.3 |
| 7 | 1 | 41650226 | T | C | SNV | intergenic_region | 0.4 |
| 8 | 2 | 333872 | C | G | SNV | intergenic_region | 0.3 |
| 9 | 2 | 1275375 | A | G | SNV | missense_variant | 0.3 |
| 10 | 2 | 4058285 | A | G | SNV | missense_variant | 0.3 |
| 11 | 2 | 27324958 | T | G | SNV | synonymous_variant | 0.3 |
| 12 | 2 | 35303751 | C | G | SNV | missense_variant | 0.3 |
| 13 | 3 | 2291086 | C | T | SNV | upstream_gene_variant | 0.3 |
| 14 | 3 | 7299077 | T | G | SNV | downstream_gene_variant | 0.3 |
| 15 | 3 | 7299101 | C | T | SNV | downstream_gene_variant | 0.3 |
| 16 | 3 | 7299136 | T | C | SNV | downstream_gene_variant | 0.3 |
| 17 | 3 | 16117473 | A | G | SNV | missense_variant | 0.3 |
| 18 | 3 | 18126184 | G | A | SNV | upstream_gene_variant | 0.3 |
| 19 | 3 | 18126206 | C | T | SNV | upstream_gene_variant | 0.3 |
| 20 | 3 | 18126212 | T | G | SNV | upstream_gene_variant | 0.3 |
| 21 | 3 | 21977903 | T | C | SNV | intergenic_region | 0.3 |
| 22 | 3 | 24394084 | G | C | SNV | intergenic_region | 0.6 |
| 23 | 3 | 31563509 | T | G | SNV | upstream_gene_variant | 0.4 |
| 24 | 3 | 32587991 | A | G | SNV | 3_prime_UTR_variant | 0.3 |
| 25 | 3 | 33876946 | T | C | SNV | intergenic_region | 0.3 |
| 26 | 4 | 2001692 | A | G | SNV | upstream_gene_variant | 0.5 |
| 27 | 4 | 2660998 | T | C | SNV | intergenic_region | 0.2 |
| 28 | 4 | 24478866 | A | G | SNV | missense_variant | 0.4 |
| 29 | 4 | 26122389 | A | G | SNV | upstream_gene_variant | 0.3 |
| 30 | 5 | 1441372 | T | G | SNV | missense_variant | 0.3 |
| 31 | 5 | 22346870 | T | G | SNV | intergenic_region | 0.3 |
| 32 | 6 | 2633791 | A | G | SNV | synonymous_variant | 0.3 |
| 33 | 6 | 11572805 | C | A | SNV | upstream_gene_variant | 0.4 |
| 34 | 6 | 22161186 | G | A | SNV | downstream_gene_variant | 0.2 |
| 35 | 7 | 2261623 | C | G | SNV | downstream_gene_variant | 0.3 |
| 36 | 7 | 8361152 | T | G | SNV | upstream_gene_variant | 0.3 |
| 37 | 7 | 10559536 | G | A | SNV | intergenic_region | 0.9 |
| 38 | 7 | 12512322 | T | G | SNV | intergenic_region | 0.3 |
| 39 | 7 | 14925810 | G | T | SNV | intergenic_region | 0.4 |
| 40 | 7 | 16205762 | A | G | SNV | 5_prime_UTR_variant | 0.4 |
| 41 | 7 | 17868859 | G | A | SNV | upstream_gene_variant | 0.3 |
| 42 | 7 | 21338148 | T | C | SNV | upstream_gene_variant | 0.4 |
| 43 | 7 | 27268351 | G | A | SNV | intergenic_region | 0.3 |
| 44 | 7 | 27268374 | C | T | SNV | intergenic_region | 0.3 |
| 45 | 7 | 27268384 | C | T | SNV | intergenic_region | 0.3 |
| 46 | 7 | 27268389 | G | A | SNV | intergenic_region | 0.3 |
| 47 | 7 | 28775765 | T | C | SNV | upstream_gene_variant | 0.4 |
| 48 | 8 | 385260 | A | G | SNV | upstream_gene_variant | 0.3 |
| 49 | 8 | 1008082 | T | G | SNV | missense_variant | 0.4 |
| 50 | 8 | 7385170 | A | C | SNV | intergenic_region | 0.4 |
| 51 | 8 | 10517046 | G | C | SNV | intergenic_region | 0.3 |
| 52 | 8 | 23725717 | A | G | SNV | missense_variant | 0.3 |
| 53 | 9 | 7615565 | T | C | SNV | intergenic_region | 0.3 |
| 54 | 9 | 11890711 | G | A | SNV | intron_variant | 0.3 |
| 55 | 9 | 16375513 | C | T | SNV | intergenic_region | 0.3 |
| 56 | 9 | 21276706 | A | C | SNV | missense_variant | 0.3 |
| 57 | 10 | 6965329 | G | A | SNV | intergenic_region | 0.3 |
| 58 | 11 | 2789079 | A | G | SNV | missense_variant | 0.3 |
| 59 | 11 | 6383147 | A | G | SNV | missense_variant | 0.3 |
| 60 | 11 | 7332284 | C | G | SNV | 3_prime_UTR_variant | 0.4 |
| 61 | 11 | 24929146 | A | G | SNV | upstream_gene_variant | 0.3 |
| 62 | 12 | 5078523 | A | G | SNV | intron_variant | 0.3 |
| 63 | 12 | 6811495 | C | G | SNV | missense_variant | 0.3 |

**Supplemental Table 4.** Number of mutation output from mutation analysis pipeline and those confirmed using IGV

| | SNV | INS | DEL | Total |
|---|---|---|---|---|
| Numbers of mutation output from mutation analysis pipeline (a) | 125 | 1 | 0 | 126 |
| Numbers of mutation confirmed using IGV (b) | 63 | 0 | 0 | 63 |
| Accuracy rate (b/a × 100) | 50.4 | 0 | - | 50.0 |
